# Herboxidiene and EGFR expression in A172 glioblastoma cells

**DOI:** 10.64898/2026.11.02.762382

**Authors:** Seiji Naito, Yuta Horita, Hiroyoshi Harada, Masayoshi Mori, Yusuke Murata, Kenji Ohe

## Abstract

Glioblastoma carries a five-year survival of about 10%, and EGFR amplification is one of the molecular criteria that define the disease in the 2021 WHO classification of central nervous system tumours. SF3b inhibitors, which displace the U2 snRNP from the intron branch point, suppress EGFR expression in lung adenocarcinoma cells by causing retention of EGFR intron 2. We asked whether the same route operates in glioblastoma, using A172 cells, which express wild-type EGFR together with an in-frame tandem duplication of exons 18–26. Herboxidiene inhibited A172 proliferation: DMSO-treated cultures reached 2.32 ± 0.61 times their day 0 reading by day 3, whereas every herboxidiene concentration from 0.125 to 1 µM held the cultures at or below their starting value and differed from the vehicle on days 2 and 3 (Dunnett, p < 0.05). RT-PCR across EGFR exons 2–3 showed no intron 2-retained product at any concentration; instead the spliced product decreased relative to ACTB, to 0.43 of the vehicle value at 0.125 µM, without deepening further up to 1 µM. EGFR protein fell, but the two forms behaved differently: the 170 kDa wild-type band was no longer detectable from 0.13 µM, whereas the 190 kDa duplicated form persisted at every concentration and declined only gradually, to 0.30 of the vehicle value at 1 µM. RT-PCR of CDKN1B (p27) showed retention of introns 1 and 2 at 0.5 µM in A172 and in A549 lung adenocarcinoma cells. In A172 cells, therefore, herboxidiene suppresses EGFR expression without the intron 2 retention seen in lung adenocarcinoma lines, and the growth inhibition is not accounted for by loss of EGFR alone.

The proliferation data are the mean ± SD of three series; the RT-PCR and western blot experiments were each performed once, without replicates or statistical testing, and should be read as preliminary.

## Introduction

Glioblastoma is a WHO grade IV tumour of the central nervous system with a five-year survival of about 10%. Surgery followed by radiotherapy with temozolomide remains the standard of care; among drugs, only temozolomide carries a strong recommendation in the Japanese guideline for adult glioblastoma, while nimustine, carmustine and bevacizumab are recommended weakly. Oncolytic herpes virus therapy (G47Δ) has entered clinical use in Japan, but no standard regimen exists after recurrence.

In the fifth edition of the WHO classification of central nervous system tumours (2021), an IDH-wildtype diffuse astrocytic glioma is classified as glioblastoma when it carries any of TERT promoter mutation, chromosome 7 gain with chromosome 10 loss, or EGFR amplification, even in the absence of the histological features. EGFR is therefore a defining molecular feature of the disease, and reducing EGFR expression is a plausible therapeutic aim.

Herboxidiene (GEX1A) binds SF3B1 and blocks the recognition of the intron branch-point adenosine by the U2 snRNP–SF3B complex, which leaves introns unspliced. SF3B inhibitors have been developed as candidates for myelodysplastic syndromes and have shown activity against solid tumours. In non-small-cell lung cancer cell lines, we have previously found that herboxidiene suppresses EGFR expression with retention of EGFR intron 2, irrespective of EGFR mutation status (Fukagawa et al., 2026). Whether this mechanism extends to glioblastoma has not been examined.

A further consideration for brain tumours is delivery across the blood–brain barrier. Herboxidiene is water-soluble but of low molecular weight, and a related study in a Niemann–Pick type C1 model reported central nervous system activity, which makes it the more tractable SF3B inhibitor to carry into an in vivo glioblastoma model if an effect is seen in vitro.

Here we compare the response of A172 glioblastoma cells with that of A549 lung adenocarcinoma cells. A172 expresses wild-type EGFR (170 kDa) together with an in-frame tandem duplication of exons 18–26 that duplicates the tyrosine kinase and calcium internalisation domains (190 kDa) and confers ligand-independent growth; A549 carries no EGFR mutation but expresses the receptor at high level. We examined proliferation, EGFR mRNA and protein, and the splicing of CDKN1B (p27).

## Materials and methods

### Reagents and antibodies

Herboxidiene (FOCUS Biomolecules) was dissolved in DMSO. Antibodies were anti-EGFR (4267T, Cell Signaling Technology), anti-ACTB (A00702, GenScript) and anti-mouse IgG HRP-linked (7076S, Cell Signaling Technology).

### Cell culture

A172 glioblastoma cells were maintained in DMEM (Nacalai Tesque) with 10% FBS (Japan Bio Serum) at 37°C under 5% CO_2_. A172 carries an in-frame tandem duplication of EGFR exons 18–26, which duplicates the tyrosine kinase and calcium internalisation domains and yields a 190 kDa receptor alongside the 170 kDa wild-type protein, and grows independently of EGF. A549 lung adenocarcinoma cells, EGFR wild-type and maintained under the same conditions, were used only in the CDKN1B experiment shown in Figure 4.

### Proliferation assay

Cells were seeded in 24-well plates at 2 × 10^4^ per well. Herboxidiene was added at 0.125, 0.25, 0.5 and 1 µM, with DMSO as the vehicle control, and the cultures were followed for 72 h. At 24, 48 and 72 h the cells were stained with crystal violet (Crystal Violet Assay Kit, Abcam) and the absorbance was read on a microplate reader (Tecan). Values are expressed as the ratio to the absorbance at 0 h of the same series.

The assay was run as three series. The first was begun on 5 August 2022 with the cells in RPMI 1640, and the second and third on 27 August 2022 in DMEM as two separate plates seeded on the same day. The three series are therefore two independent experiments, one of them performed in duplicate plates, and the medium of the first differs from the other two. Absorbance was read at 570 nm for these series rather than at 595 nm.

### RT-PCR

Cells were seeded in 6-well plates at 5 × 10^5^ per well, treated with herboxidiene at the concentrations used in the proliferation assay, and harvested after 24 h. Total RNA was extracted with Trizol (Invitrogen): after chloroform extraction, RNA was recovered from the aqueous phase. One microgram of total RNA was reverse-transcribed with ReverTra Ace (TOYOBO), and PCR was performed with Ex Taq (TaKaRa). The primers are listed below. Products were resolved on agarose gels and visualized with ethidium bromide.

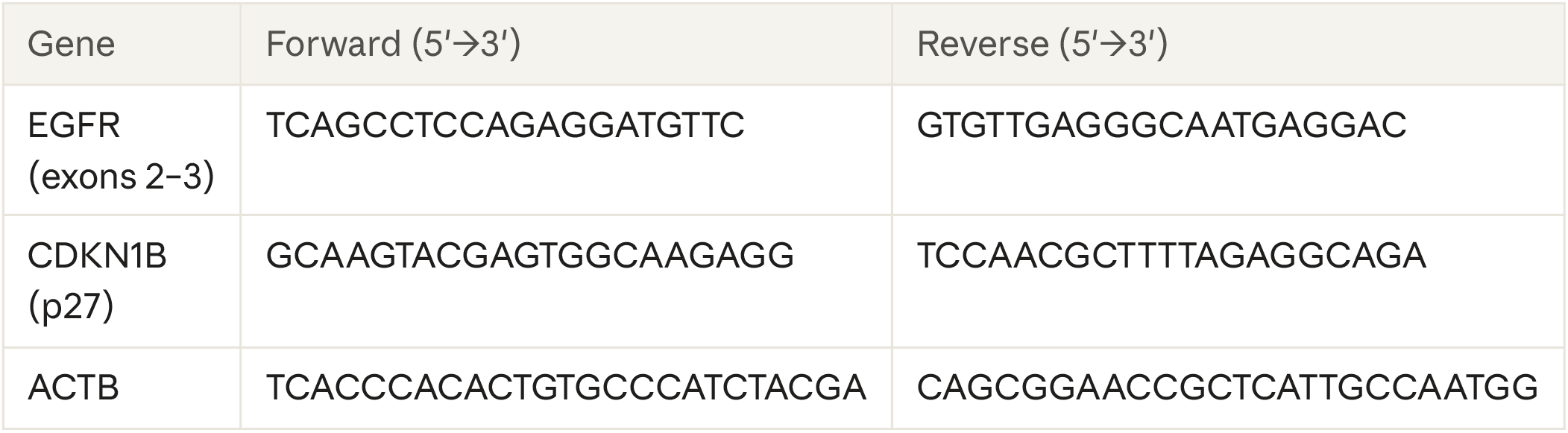

### Western blotting

Protein was recovered by acetone precipitation from the interphase remaining after the Trizol/chloroform extraction used for RNA. Pellets were dissolved in urea sample buffer (10 mM Tris pH 7.5, 25 mM KCl, 5 mM MgCl_2_, 6 M urea) and sonicated to remove co-precipitated DNA and RNA. Protein was quantified with Pierce 660 nm Protein Assay (Thermo Fisher) and approximately 20 µg per lane was separated by SDS-PAGE and transferred to PVDF membrane by tank blotting. Membranes were blocked with Blocking One (Nacalai Tesque) for 10 min at 4°C, incubated with primary antibody (1:1000) for 16 h at 4°C and with secondary antibody (1:5000) for 1 h at room temperature, and developed with Chemi-Lumi One L (Nacalai Tesque).

### Replication and analysis

Proliferation values are the mean ± SD of the three series described above, compared with the DMSO control at each time point by Dunnett’s test (scipy.stats.dunnett), which corrects for the four concentrations tested against one control. The RT-PCR and western blot experiments were each performed once; band intensities in Figures 2 and 3 were quantified densitometrically and normalised to ACTB, and these ratios are descriptive with no statistical test applied.

The A549 proliferation, EGFR RT-PCR and EGFR western blot experiments performed alongside this work are reported separately (Fukagawa et al., 2026) and are not reproduced here. Figure 4 is the only panel in this manuscript that includes A549.

## Results

### Herboxidiene inhibits proliferation of A172 cells

A172 cells were followed for three days after the addition of herboxidiene (Figure 1). Pooling the three series, DMSO-treated cultures reached 2.32 ± 0.61 times their day 0 reading by day 3. Herboxidiene held the cultures at or below their starting value: the day 3 ratios were 0.74 ± 0.19 at 0.125 µM, 0.77 ± 0.23 at 0.25 µM, 0.73 ± 0.22 at 0.5 µM and 0.76 ± 0.23 at 1 µM. By Dunnett’s test against DMSO, all four concentrations differed from the vehicle on day 2 (p = 0.041, 0.021, 0.018 and 0.018) and on day 3 (p = 0.0006, 0.0008, 0.0007 and 0.0006). None differed on day 1 (p = 0.39–0.63).

**Figure 1.**
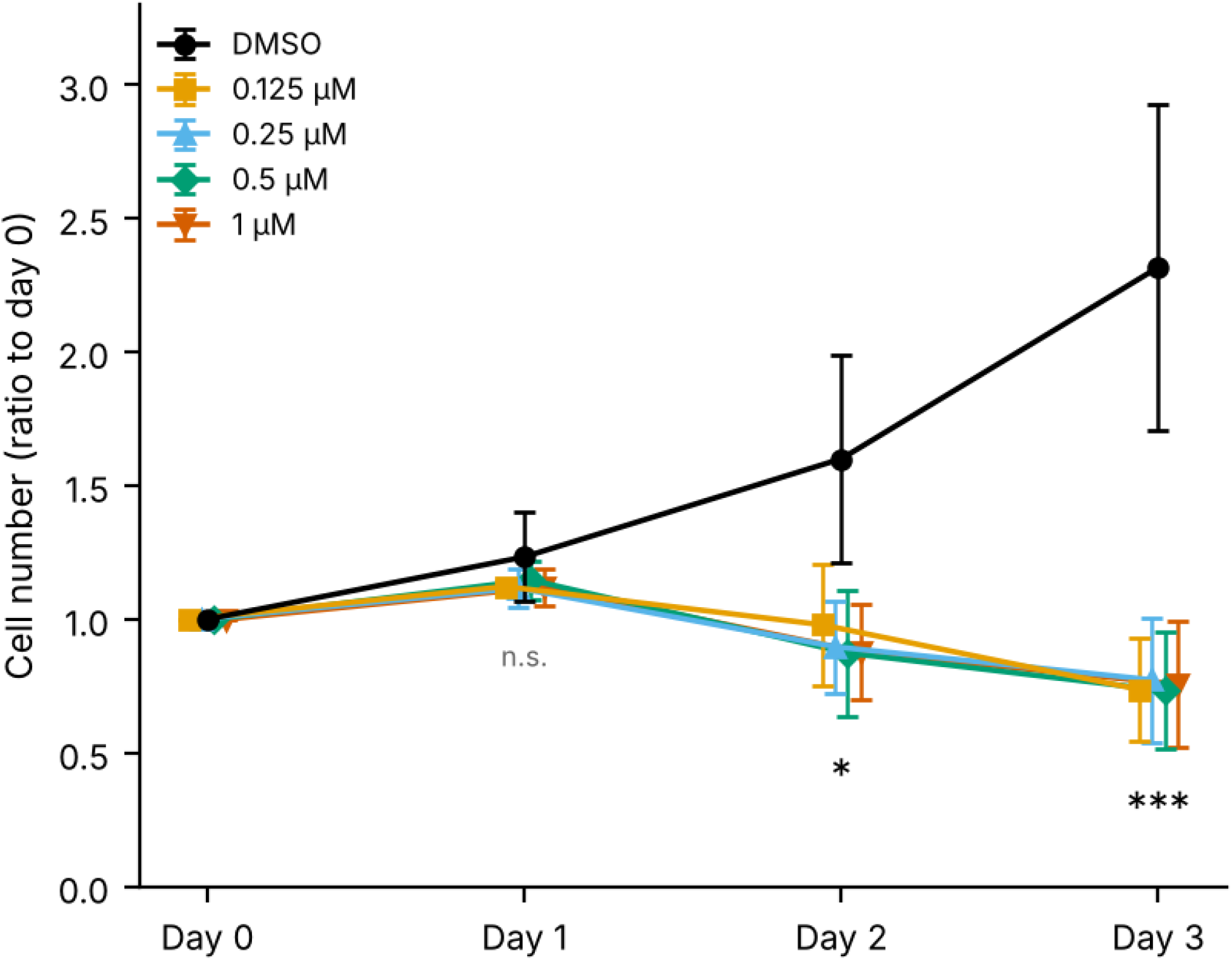
Herboxidiene inhibits proliferation of A172 cells. Cells were seeded at 2 × 10^4^ per well in 24-well plates and treated with herboxidiene at 0.125, 0.25, 0.5 and 1 µM or with DMSO. Viable cells were stained with crystal violet at 24, 48 and 72 h and the absorbance is expressed as the ratio to the 0 h reading of the same series. Points are the mean of three series and bars are the SD; the three series are two independent experiments, one of them run as duplicate plates, and the first was carried out in RPMI 1640 rather than DMEM. Asterisks give the least significant of the four Dunnett comparisons against DMSO at that time point (* p < 0.05, *** p < 0.001); n.s., no concentration differed from DMSO.

The mean cell number fell between day 2 and day 3 at every concentration, so the cultures lost cells rather than merely stopping their growth. The four concentrations were not distinguishable from one another at any time point, so no concentration–response relationship was resolved over the range tested: 0.125 µM, the lowest concentration used, already produced the full effect, and the potency of herboxidiene in this line is therefore bounded rather than determined.

### EGFR mRNA falls without retention of intron 2

RT-PCR across EGFR exons 2–3 was performed 24 h after treatment (Figure 2). The primers used here give a spliced product of 139 bp and an intron 2-retained product of 1006 bp, and in lung adenocarcinoma lines the retained product appears on treatment and increases with concentration. In A172, no retained product was detected at any concentration. The spliced product instead decreased relative to ACTB: 1.0 in untreated cells, 0.85 under DMSO and 0.35 at 0.125 µM (Figure 2A). Across the concentration series the EGFR/ACTB ratio, expressed relative to DMSO, was 0.43 at 0.125 µM, 0.26 at 0.25 µM, 0.34 at 0.5 µM and 0.41 at 1 µM (Figure 2B). The reduction was therefore present from the lowest concentration tested but did not deepen above 0.25 µM.

**Figure 2.**
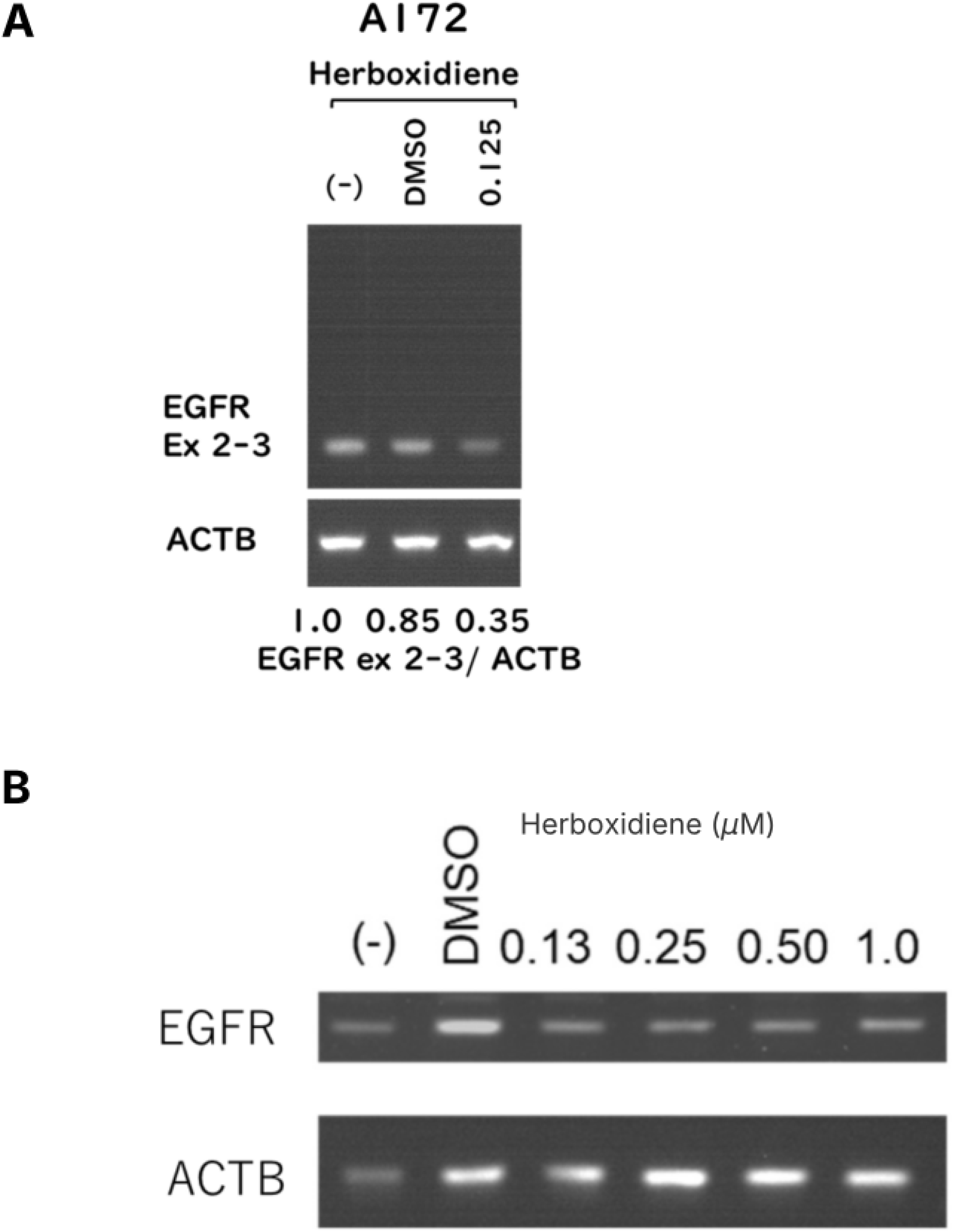
EGFR mRNA falls in A172 cells without retention of intron 2. RT-PCR across EGFR exons 2–3 after 24 h of treatment; the primers give a spliced product of 139 bp and an intron 2-retained product of 1006 bp. (A) Untreated, DMSO and 0.125 µM herboxidiene, with ACTB below. The EGFR/ACTB ratios shown beneath the panel are 1.0, 0.85 and 0.35. (B) The full concentration series: untreated, DMSO, and herboxidiene at 0.13, 0.25, 0.50 and 1.0 µM, with ACTB below; EGFR/ACTB ratios relative to DMSO are 0.86, 1.00, 0.43, 0.26, 0.34 and 0.41. No intron 2-retained product was detected in any lane. Single experiment.

### The two EGFR forms in A172 respond differently

EGFR protein was examined 24 h after treatment (Figure 3). Untreated and DMSO lanes showed two bands: the 190 kDa product of the exon 18–26 tandem duplication and the 170 kDa wild-type receptor. The 170 kDa band was no longer detectable from 0.13 µM upwards. The 190 kDa band persisted in every treated lane and declined gradually, the EGFR/ACTB ratio relative to DMSO falling from 1.02 at 0.13 µM to 0.92 at 0.25 µM, 0.67 at 0.5 µM and 0.30 at 1 µM. ACTB was unchanged. The concentration at which the wild-type band disappeared, 0.13 µM, is the same concentration at which the mRNA reduction was already near its maximum, whereas the duplicated form was still present at concentrations that caused loss of cells.

**Figure 3.**
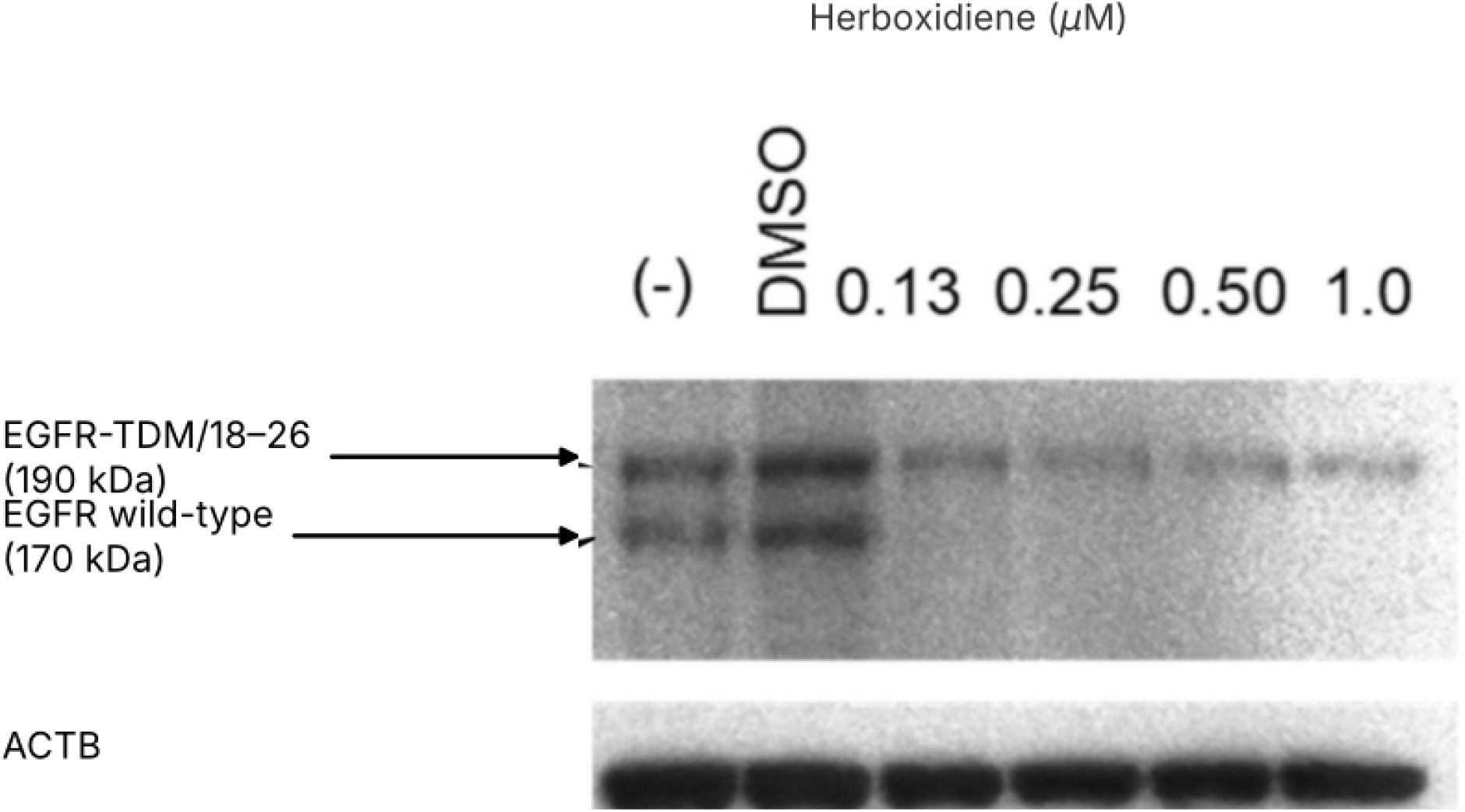
The 170 kDa and 190 kDa EGFR species respond differently to herboxidiene. Western blot for EGFR (C-terminal antibody 4267T) and ACTB after 24 h of treatment: untreated, DMSO, and herboxidiene at 0.13, 0.25, 0.50 and 1.0 µM. The upper band is the 190 kDa product of the EGFR exon 18–26 tandem duplication and the lower band the 170 kDa wild-type receptor. EGFR/ACTB ratios relative to DMSO are 1.10, 1.00, 1.02, 0.92, 0.67 and 0.30. Single experiment.

### CDKN1B (p27) introns are retained

Because growth inhibition was more marked than the loss of EGFR protein at the same concentrations, we examined CDKN1B (p27), for which SF3b inhibition has been reported to produce a truncated 22 kDa isoform. RT-PCR spanning exon 1 to exon 3 was performed on cells treated with 0.5 µM herboxidiene (Figure 4). In A172 a slower-migrating band appeared on treatment, corresponding in size to the transcript retaining introns 1 and 2, alongside the fully spliced exon 1–2–3 product seen in the mock lane. A549 cells treated in the same experiment showed the same change, of comparable extent. Unlike EGFR, therefore, CDKN1B responded to herboxidiene by intron retention in the glioblastoma line as well.

**Figure 4.**
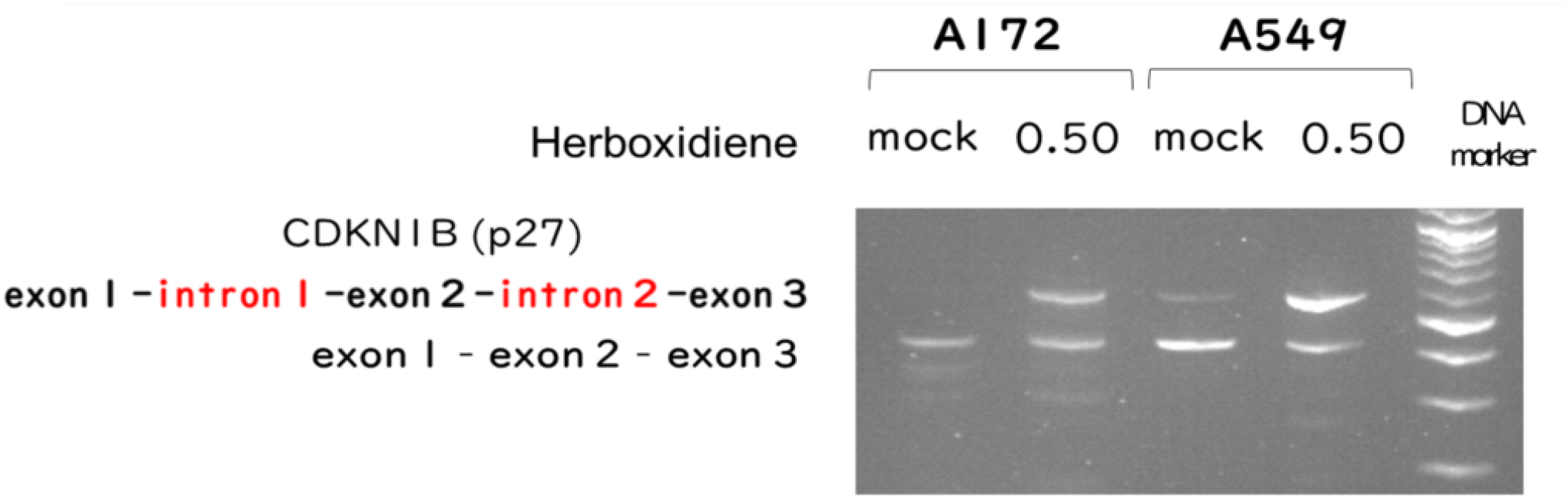
Herboxidiene induces retention of CDKN1B introns 1 and 2. RT-PCR spanning CDKN1B exon 1 to exon 3 after 24 h of treatment with 0.5 µM herboxidiene or mock, in A172 and A549 cells, with a DNA size marker in the right-hand lane. The slower-migrating band corresponds in size to the transcript retaining introns 1 and 2 and the faster band to the spliced exon 1–2–3 product; band identity was inferred from size and was not confirmed by sequencing. Single experiment.

## Discussion

The main observation is a negative one with a positive counterpart: in A172 glioblastoma cells, herboxidiene reduced EGFR mRNA and protein, but did so without producing the EGFR intron 2-retained transcript that accompanies the same reduction in lung adenocarcinoma lines. Since intron 2 retention has been used as the readout of target engagement for SF3b inhibitors at this locus, its absence in a line where EGFR nonetheless falls indicates that the fall is reached by another route.

Three routes are compatible with the data. First, retention of an EGFR intron other than intron 2, which the exon 2–3 primers would not detect. Second, suppressed splicing of a transcription factor that drives EGFR expression, so that the loss of EGFR mRNA is secondary. Third, suppressed splicing of a factor governing EGFR mRNA stability. The experiments reported here do not separate them. A reverse-transcription survey across the remaining EGFR introns would address the first directly and is the natural next step; RNA sequencing would address all three at once and would also show how much of the transcriptome is affected at these concentrations.

The behaviour of the two EGFR protein species deserves comment. The 170 kDa wild-type band disappeared from 0.13 µM, while the 190 kDa product of the exon 18–26 tandem duplication was still present at 1 µM. Since the duplication is in frame and the mRNA encoding the 170 kDa protein is contained within that encoding the 190 kDa protein, the two transcripts share their promoter, their 3’ untranslated region and all introns outside the duplicated stretch. A difference in their response is therefore not readily explained by an effect on any shared element, and we cannot account for it from these data. One possibility is that the junction created between exon 26 and exon 18 in the duplicated allele is spliced by a route that differs in its dependence on SF3b. We note it as unexplained rather than resolved.

Growth inhibition in A172 is unlikely to be attributable to loss of EGFR. The cell number fell at 0.5 and 1 µM, concentrations at which the duplicated receptor was still detectable, and published knockdown work indicates that removing EGFR does not by itself arrest glioblastoma lines. The CDKN1B result points to a broader explanation: introns 1 and 2 of CDKN1B were retained at 0.5 µM in A172, as in A549. Retention here is predicted to yield a C-terminally truncated p27 of about 22 kDa, which has been reported to accumulate after SF3b inhibition and to retain cyclin-dependent kinase inhibitory activity. Growth inhibition by herboxidiene is therefore better read as the sum of splicing changes across many transcripts, of which EGFR is one, than as an EGFR-directed effect. We did not measure p27 protein, so the link between the retained transcript and the truncated isoform is inferred, not shown.

For glioblastoma specifically, two features make SF3b inhibition worth pursuing. EGFR amplification is one of the molecular criteria that define the disease, so any agent that lowers EGFR expression addresses a defining feature rather than an incidental one. And herboxidiene, unlike most SF3b inhibitors, has been reported to act in the central nervous system in a non-oncological model, which makes a blood–brain barrier-penetrant candidate conceivable. Neither point is established for the tumour setting, and an orthotopic model would be needed before either could be claimed.

### Limitations

The proliferation assay rests on three series, but these are two independent experiments rather than three, since two of the series were plates seeded on the same day, and the first series was carried out in RPMI 1640 while the other two were in DMEM. The Dunnett comparisons treat the three as independent, which overstates the degrees of freedom; the day 3 differences are large enough that this is unlikely to change the conclusion, but the day 2 p values, which sit between 0.018 and 0.041, should be read with that in mind. No concentration–response relationship was resolved, because the lowest concentration tested already produced the full effect.

The RT-PCR and western blot experiments were each performed once, so the band intensities and the ratios derived from them carry no estimate of variability and no statistical test was applied. The absence of an intron 2-retained product in A172 is a negative result from one RT-PCR; ACTB amplified normally in the same reactions, but the negative would be more secure with a positive control for retention run on the same gel. The identity of the slower-migrating CDKN1B band was inferred from its size and was not confirmed by sequencing, and p27 protein was not measured, so the link between the retained transcript and the truncated isoform is inferred rather than shown. All work was in a single glioblastoma line in vitro; whether the pattern holds in other glioblastoma lines, in cells carrying EGFRvIII rather than the exon 18–26 duplication, or in vivo, is untested.

## Declarations

### Author contributions

Conceptualization: K.O. Methodology: K.O. Investigation: S.N. and K.O. Formal analysis: K.O. Visualization: K.O. Writing — original draft: K.O. Supervision and project administration: K.O. Y.H., H.H., M.M. and Y.M. advised on study design and reviewed the manuscript.

### Data availability

The data supporting the findings are contained within the figures. Original gel and blot images are available from the corresponding author on reasonable request.

### Related work

The A549 proliferation, EGFR RT-PCR and EGFR western blot experiments performed in the same laboratory with the same compounds are reported in Fukagawa et al. (2026) and are not reproduced here.

